# Falign: An effective alignment tool for long noisy 3C data

**DOI:** 10.1101/2022.10.30.514399

**Authors:** Ying Chen, Zhuo-Bin Lin, Long-Jian Niu, Jia-Yong Zhong, Yi-Zhi Liu, Chun-Hui Hou, Feng Luo, Chuan-Le Xiao

## Abstract

Fragmented long noisy reads (FLNRs), such as Pore-C, contain multiple fragments of varied length separated by restriction enzyme sites. Existing alignment tools have a low mapping rate for short fragments and find incorrect fragment boundaries, which affects the utilization of FLNRs for downstream studies. Here, we develop Falign, a sequence alignment method that is adapted to the nature of FLNRs. Falign adopts a two-phase approach to efficiently align both long and short fragments. Falign uses the restriction enzyme sites on the reference genome as boundaries, which avoids the problem of destroyed fragment boundaries on FLNRs. Falign employs a multiple-stage searching mechanism to effectively recover the alignments of FLNRs with multiple fragments and interchromosomal fragments. Experiments on simulated and experimental fragmented long noisy 3C datasets show that Falign can effectively recover the constructs of reads and the sampled loci of the fragments. Falign allows significantly higher data utilization for FLNRs.

## Introduction

Three-dimensional (3D) chromatin organization plays an important role in transcriptional programs and is essential in cell epigenetic regulation^1^. In the last decade, sequencing-based chromosome conformation capture (3C) technologies, including second-order 3C^2^, 4C^3,4^, 5C^5^, ChIA-PET^6^ and Hi-C^7^, and high-order SPRITE^8^ and GAM^9^, have been developed to investigate the 3D structure of chromosomes inside the nucleus by measuring the contact frequencies between two or more chromatin fragments. 3C technologies have been successfully used in studies of genomic 3D architecture^10^, de novo genome assembly^11-13^, gene regulation^14,15^, and DNA replication timing^16^. Recently, technologies such as Pore-C^17,18^ and C-walk that couple 3C with long-read sequencing were developed. Compared to previous 3C technologies, these new 3C technologies have advantages in identifying high-order 3D structures in genome loci^17,18^ due to their ability to sequence multiple concatemers using long reads.

A Pore-C read usually contains several nonoverlapping chromatin fragments with lengths from tens of bps to thousands of bps. The fragments are separated by restriction enzyme sites. However, inside each fragment, there can also exist one or more restriction enzyme sites. Similar to other 3C assays, the first step of analyzing Pore-C reads is to align the Pore-C reads to the corresponding reference genome to identify interacting fragments encoded in them. However, the number and length of fragments in a Pore-C read are undetermined *a priori*. Therefore, mapping Pore-C reads to the reference genome includes two challenges: correctly decomposing the read into fragments and mapping the fragments to the correct loci of the reference genome.

Currently, bwa^19^ is used to align Pore-C reads to loci in the genome^17^. NGMLR^20^ and minimap2^21^ are two widely used alignment tools that were designed for aligning noisy long reads. However, the fragmented nature of Pore-C reads makes their alignment difficult. All these tools suffer from two drawbacks when they are applied to align Pore-C reads (see Results). First, the mapping rates for short fragments are low. The loss of short fragments leads to over 50% of Pore-C reads failing to be completely mapped (there is no alignment result for at least one fragment in the read). Second, there are many incorrect fragment boundaries among the alignment results. Incorrect fragment boundaries not only make the decomposition of Pore-C reads incorrect but also lead to incorrect results in subsequent analyses, such as base methylation modification detection in fragment boundaries. Furthermore, the loss or incorrect mapping of one fragment will lead to the loss of N-1 pairwise contacts, where N is the total number of fragments in a read. Overall, these two drawbacks can result in a loss of 20-80% of pairwise contacts, which significantly reduces the effective utilization of Pore-C data.

In this paper, we develop Falign, an alignment tool that adapts to fragmented long noisy reads, such as Pore-C reads. Falign takes a two-phase approach. In the first phase, we develop a local DDF chain scoring algorithm^22^ to select fragment candidates and then carefully extend the long fragment candidates by considering restriction enzyme sites as boundaries during further local alignment. In the second phase, we select short fragments and develop a dynamic programming-based method to generate the most plausible set of fragment alignments as the mapping result. Experimental results on simulated and experimental datasets show that Falign solves the short fragment and incorrect boundary problems, is much faster than other aligners and achieves significantly higher data utilization. We also demonstrate that Falign obtains good performance on other fragmented long noisy 3C datasets, such as C-walks^23^ and MC-3C^24^.

## Results

### Challenges in aligning Pore-C reads

To understand the bottlenecks of current methods for aligning Pore-C reads, we performed a simulation study using bwa, NGMLR and minimap2. We first simulated Pore-C datasets of *H. sapiens, A. thaliana* and *D. melanogaster* based on the DpnII restriction enzyme (Methods and Supplementary Note 1). We sampled various numbers of fragments with various lengths from restriction sites on the reference genome and concatenated them into one sequence. Then, we modified DeepSimulator^25^ to directly generate errors mimicking Oxford Nanopore Technologies (ONT) sequencing based on the concatenated sequence. Read information for the three datasets is shown in Supplementary Table 1. The average lengths of fragments in the three datasets were 873, 844, and 898, with the length of the majority of fragments being 200-1200 bp (Supplementary Figure 1A). The sequencing error rates were 8% on average and mainly distributed between 6% and 12% (Supplementary Figure 1B). The proportions of reads composing 1, 2, 3, 4, 5 and ≥ 6 fragments were 5.66%, 5.37%, 8.26%, 10.03%, 11.02% and 59.66%, respectively (Supplementary Figure 1C). The proportions of reads whose fragments were spread across 1, 2, 3, and ≥ 4 chromosomes were 38.60%, 34.00%, 16.63% and 10.77%, respectively (Supplementary Figure 1D).

Then, we ran bwa, NGMLR and minimap2 on these datasets (Methods and Supplementary Notes 2-3). As shown in Figure 1 A, for the *H. sapiens* dataset, we found that there were a large number of fragments (34.45%, 44.25% and 17.69%) that failed to be mapped back to the reference genome by bwa, NGMLR and minimap2. Most of the unmapped fragments were shorter than 1000 bp (Figure 1B). The shorter a fragment is, the more likely its alignment is to be missing. Consequently, 78.01%, 85.85% and 91.84% of simulated Pore-C reads were partially mapped in the results of bwa, NGMLR and minimap2, respectively (Figure 1 C). Furthermore, the percentage of completely mapped reads decreased as the number of fragments in the reads increased (Figure 1D). For reads with more than 6 fragments, only 2.33%, 3.42% and 22.32% reads were completed and mapped by bwa, NGMLR and minimap2, respectively (Figure 1D). Among the correctly mapped fragments, there were 25.99%, 48.46% and 56.59% overlapped fragments (fragments having large overlaps (> 50 bp) with adjacent fragments) in the results of bwa, NGMLR and minimap2, respectively, which led to 51.47%, 69.92% and 85.19% of reads containing overlapped fragments in the results of bwa, NGMLR and minimap2, respectively (Figure 1E). These results show that there were a significant number of fragments that were either not mapped or had incorrectly resolved boundaries in the alignment results. The mapping results of the simulated *A. thaliana* and *D. melanogaster* datasets were also similar (Supplementary Figure 2 and Table 1). Furthermore, the low mapping rate of fragments led to the loss of a large number of pairwise contacts among the multiway contacts, which are the basic units of genomic 3D architecture. For example, bwa, NGMLR and minimap2 lost approximately 57%, 65% and 32% of the pairwise contacts, respectively, and the precision of their pairwise contacts was more than 90% on the H. sapiens dataset (Figure 1F-G). The loss of a significant amount of pairwise contacts could lead to an incomplete and even biased reconstruction of genomic 3D architecture.

**Figure 1.**
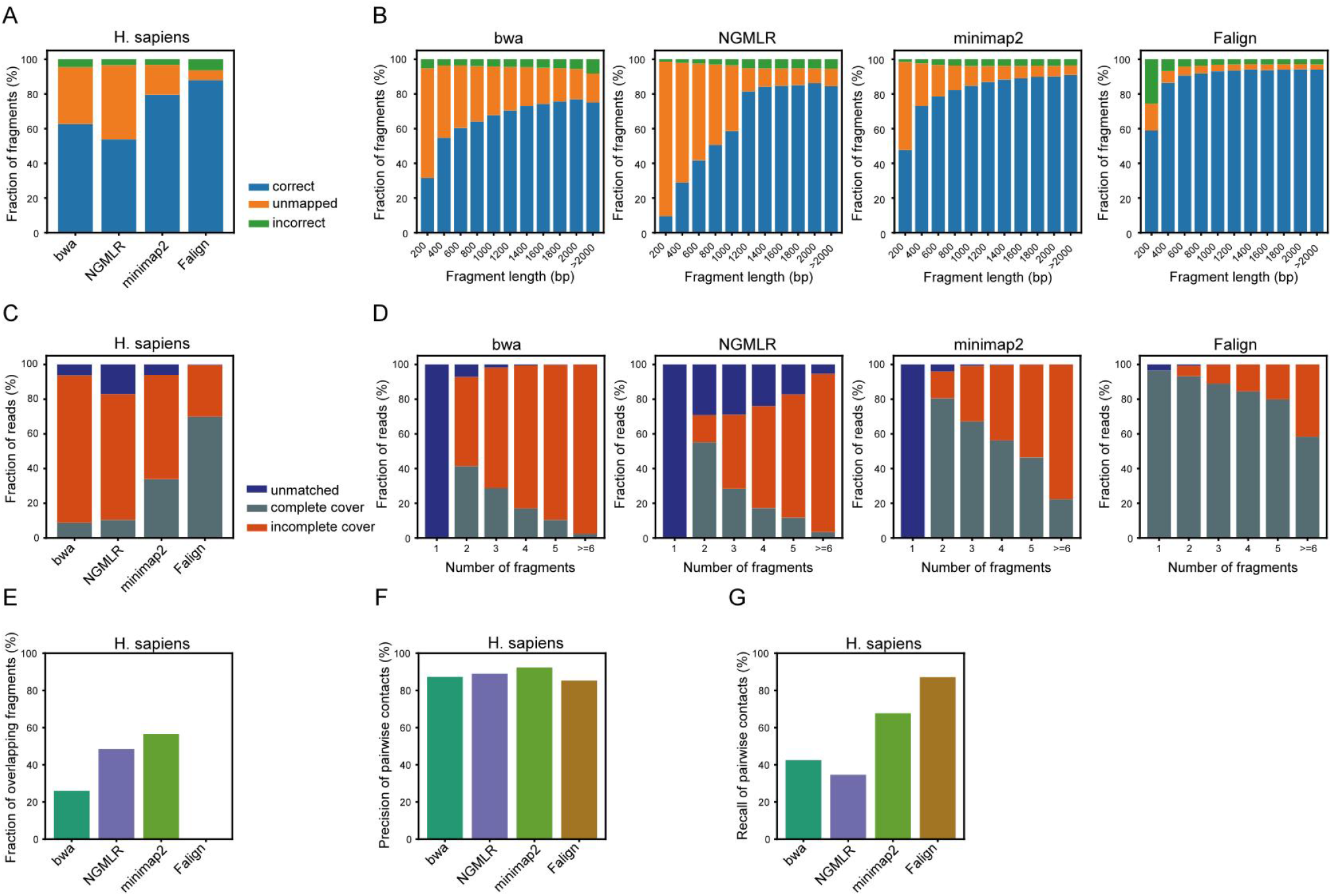
Mapping results on the simulated Pore-C datasets. (A-B) Mapping results in terms of fragments. (C-D) Mapping results in terms of reads. (E) Fractions of overlapping fragments. (F) Precision of pairwise contacts. (G) Recall rates of pairwise contacts.

**Table 1.** Performance on simulated Pore-C datasets. The datasets were generated using the DpnII restriction enzyme. Let rf, mf, and cf denote the number of fragments in the simulated datasets, the number of fragments identified by the mapping results, and the number of correctly mapped fragments, respectively. Then, precision=100.0*cf/mf, and sensitivity=100.0*cf/rf. A fragment is called confident if one of its alignment’s four offsets (read start, read end, reference start, reference end) is located at a restriction enzyme site; it is called overlapping if it shares an overlap greater than 50 bp with its adjacent fragment. A Pore-C read is called completely mapped if almost all of its bases are mapped to the reference (see main text) and is called overlapping if it contains overlapping fragments. Map AFPR: average number of fragments per read computed from the mapping results. Raw AFPR: average number of fragments per read computed from the simulated Pore-C reads. Map contacts: number of pairwise contacts extracted from the mapping results. Correct contacts: number of pairwise contacts formed by correctly mapped fragments. Raw contacts: number of true pairwise contacts encoded in the simulated reads.

| Dataset | Aligner | Size (G)/<br>Time (h) | Precision<br>/Sensitivity (%) | Fragment (%)<br>Confident/<br>Overlapping | Read (%)<br>Complete/<br>Overlapping | AFPR<br>map<br>/raw | Contacts (M)<br>map/correct/raw |
| --- | --- | --- | --- | --- | --- | --- | --- |
| H. sapiens | bwa | 2.02/4.78 | 93.31/65.55 | 88.99/25.99 | 21.79/51.47 | 5.51/7.84 | 6.22/5.43/12.77 |
|  | NGMLR | 2.02/1.21 | 93.93/55.75 | 88.02/48.46 | 14.15/69.92 | 4.66/7.84 | 4.97/4.43/12.77 |
|  | minimap2 | 2.02/0.83 | 95.99/82.31 | 90.18/56.59 | 8.16/85.19 | 6.73/7.84 | 9.38/8.65/12.77 |
|  | Falign | 2.02/0.33 | 94.32/94.22 | 99.59/0.00 | 95.28/0.00 | 7.84/7.84 | 12.72/11.26/12.77 |
|  | Falign_ngf | 2.02/0.24 | 94.85/89.07 | 99.84/0.00 | 83.24/0.00 | 7.37/7.84 | 10.69/9.64/12.77 |
| A. thaliana | bwa | 2.28/1.43 | 93.75/66.65 | 71.21/25.13 | 22.67/50.44 | 5.58/7.85 | 6.53/5.77/13.12 |
|  | NGMLR | 2.28/0.60 | 95.43/57.03 | 69.47/48.46 | 14.66/69.81 | 4.69/7.85 | 5.19/4.76/13.12 |
|  | minimap2 | 2.28/0.54 | 97.50/84.38 | 72.64/56.46 | 8.59/85.17 | 6.79/7.85 | 9.76/9.30/13.12 |
|  | Falign | 2.02/0.08 | 96.27/95.59 | 99.98/0.00 | 98.02/0.00 | 7.79/7.85 | 12.91/11.90/13.12 |
|  | Falign_ngf | 2.02/0.07 | 96.23/92.05 | 99.97/0.00 | 90.55/0.00 | 7.51/7.85 | 11.62/10.72/13.12 |
| D. melanogaster | bwa | 2.05/1.54 | 91.18/61.30 | 84.89/23.26 | 26.25/43.94 | 5.01/7.45 | 5.54/4.64/12.40 |
|  | NGMLR | 2.05/0.57 | 94.21/48.50 | 84.72/44.72 | 17.79/62.81 | 3.83/7.45 | 3.76/3.37/12.40 |
|  | minimap2 | 2.05/0.52 | 97.01/77.79 | 87.32/51.35 | 11.73/77.13 | 5.97/7.45 | 7.97/7.53/12.40 |
|  | Falign | 2.05/0.11 | 94.12/93.45 | 99.98/0.00 | 98.26/0.00 | 7.40/7.45 | 12.20/10.85/12.40 |
|  | Falign_ngf | 2.05/0.08 | 94.59/87.59 | 99.97/0.00 | 88.10/0.00 | 6.91/7.45 | 10.23/9.27/12.40 |

Our analysis shows that current mapping methods have low mapping rates for short fragments in Pore-C reads. Since bwa is designed for mapping short and accurate next-generation reads, the low mapping rates of bwa may be due to the high error rate in the Nanopore sequencing reads. Additionally, bwa is slow in aligning long noisy reads. It takes approximately 4-5 days for bwa to align the reads of one flow cell. On the other hand, NGMLR and minimap2 are designed for noisy long reads and become less effective for short fragments. Furthermore, all current mapping methods have a large number of incorrect mapping results due to overlapped boundaries of fragment alignments. This is because none of the current methods consider the restriction enzyme sites, which are boundaries that separate fragments, during the local alignment of fragments. Therefore, it is necessary to design a new mapping tool for fragmented long noisy reads, such as Pore-C, to enable its efficient utilization.

### Overview of Falign

To address the two issues mentioned above, we developed Falign, an alignment tool for fragmental long noisy reads (FLNRs). Falign is specifically designed for increasing the mapping rate of short fragments and solving the mapping boundary, and it helps completely map reads with multiway contacts or interchromosomal contacts. Falign contains four modules (Figure 2): 1) long fragment candidate detection; 2) monosome long fragment candidate extension; 3) monosome gap filling and alignment selection; and 4) polysomy gap filling and alignment selection. Falign outputs results in SAM or PAF format^21^, and the mapping quality score of each alignment is assigned (Methods). In the following sections, we use Pore-C reads as examples to explain Falign.

**Figure 2.**
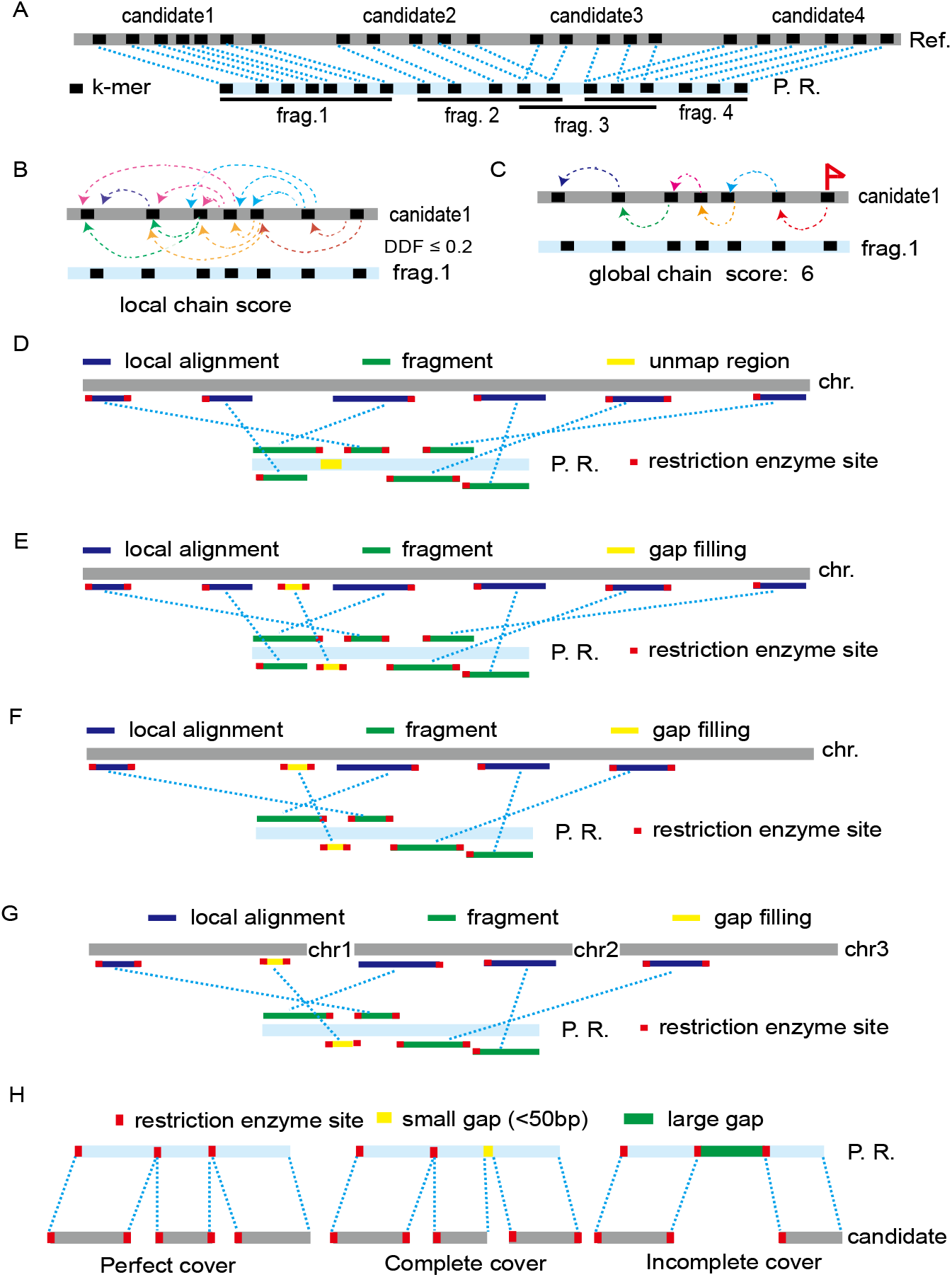
Illustration of Falign. (A) Four candidates and fragments between the read and the reference. (B-C) The detection of candidate 1 and fragment 1: B) Block chain scoring of *k* -mer matches. Two *k* -mer matches are connected by an arrow if they are close to each other and support each other by DDF. C) The *k* -mer match chain created by backtracking using the local chain scoring. This chain is considered to be a single alignment candidate with global chain score 6. (D) Confident local alignments between fragments and a chromosome. (E) Filling a gap using an alignment (yellow) between the gap region of the read and a chromosome. (F) A perfect cover consisting of five local alignments is selected from (E). If no perfect cover is found after extending all seed chromosome alignment candidates, we proceed to (D). The local alignments between the read and the seed subjects. (G) A perfect cover consisting of polysomy alignment selection. (H) The three categories of mapping results.

### Candidates for long fragment detection with local chain scoring

Similar to other seed-extension alignment tools^21,22^, the first step of Falign is to select candidate sequences in the reference genome for local alignment. For each short fragment, there may be a large number of candidates introduced by repetitive subsequences^22^, and local alignment with those candidates will be very time-consuming. Here, we develop a fast chain scoring method based on the distance difference factor (DDF), which measures the connection relation between two matched *k* -mers [22] for selecting candidates for long fragments (Figure 2A-C). For each matched *k* -mer pair, Falign considers the forward 800 bp reference sequence as its chain scoring block (Methods). The matched *K*-mer is then scored only against matched k-mers in its chain scoring block (Figure 2B). For matched *K*-mer pairs with a distance on the reference genome greater than 800 bp, Falign no longer conducts direct DDF scoring between them.

Two matched k-mer pairs with a distance on the reference genome greater than 800 bp are considered to support each other only if there exist other matched k-mer pairs forming a scoring chain between them (Supplementary Figure 3). The successive chain scoring of matched *k* -mers (starting from the one with the minimum reference position) generates various lengths of k-mer chains after backtracking (Figure 2C). Every matched *k* -mer chain is assigned a global chain score, which is the highest chain score of the matched *k* -mer in the chain (Methods). The subsequence between the left- and right-most positions in the chain is treated as the covering range of the chain, which also provides evidence for the gap-filling process described below. The k-mer chains with high scores are the alignment candidates. As demonstrated in Supplementary Figure 4, the global chain scores of the alignment candidates grow linearly with fragment length, implying that the selected candidates with high global chain scores are long fragments.

### Monosome long fragment candidate extension

We noticed that over 50% of Pore-C reads have fragments that spread only on one single chromosome. Therefore, Falign first extends long fragment candidates in one chromosome at each step (Methods). The candidates belonging to each chromosome are first sorted using chain scores. The extension process begins with the candidate with the highest score. For each candidate, Falign first aligns it with the fragment using Edlib algorithm^26^-based local alignment. The Pore-C fragments have restriction sites as their boundaries, which can be valuable criteria for alignment boundary selection. However, sequencing errors and the differences between the underlying genome and the reference genome, such as SNPs, mean that many fragment alignments do not strictly start or end at restriction enzyme sites. If Falign successfully finds restriction enzyme sites within 50 bp of the first alignment offsets, it performs another global alignment on the subsequence between restriction enzyme sites (close to the alignment offsets of the first alignment) to obtain a new detail alignment; otherwise, Falign keeps the first alignment. Fixing alignment boundaries with restriction enzyme sites is critical for eliminating overlapping fragments and overlapping reads. Current aligners (minimap2, NGMLR and bwa) do not perform this step, which results in a high proportion of incorrectly resolved fragment boundaries. For each chromosome, Falign extends candidates to obtain up to 50 alignments (Figure 2D), which represent the local alignments for long fragments.

### Monosome gap filling and alignment selection

Next, Falign performs a gap-filling step to align missed short fragments (Figure 2E and Methods). Falign first detects the missing regions not covered by any of the 50 alignments. For each k-mer chain in the chromosome that has not been processed, if over 80% of the covering range of the chain is in one of the missing regions, Falign performs the same extension procedure as those performed for long fragments and updates the missing regions with that alignment. Although we perform an exhaustive search for missing regions, the computational cost is still relatively low since the missing regions are usually short.

After gap filling, Falign finds a subset of fragment alignments that make up the read. There exist three possible cases for a Pore-C read’s fragments: 1) the fragments are sampled from one single chromosome and have restriction enzyme site boundaries; 2) the fragments are sampled from multiple chromosomes and have restriction enzyme site boundaries; and 3) the fragments do not have restriction enzyme site boundaries. Here, we develop a dynamic programming-based method to find a set of fragment alignments that can cover a whole Pore-C read. A set of alignments is called a perfect cover (Figure 2H) if the following conditions hold: 1) every alignment in the set has restriction enzyme site boundaries; 2) every base of the Pore-C read is covered by one and only one alignment in the set; and 3) no two alignments in the set have overlapping read bases or overlapping reference bases. A set of alignments is called a complete cover (Figure 2H) if the following conditions hold: 1) there is no larger than a 50 bp gap that is not covered by any alignment; 2) the total number of unmapped read bases is less than 200 bp; and 3) no two alignments in the set have overlapping read bases or overlapping reference bases. A perfect cover is also a complete cover. A set of alignments is called an incomplete cover if it is not a complete cover (Figure 2H).

Given a set of local alignments (the alignments can be from a single chromosome or from multiple chromosomes), our dynamic program method first attempts to select and return a perfect cover (Methods). If no perfect cover is found, it then tries to select and return a complete cover with a loose criterion. If no complete cover is found, it finally selects and returns an incomplete cover corresponding to a set of nonoverlapping fragments that maximally cover the read with the looser criterion. The perfect cover has the highest confidence, the complete cover is second, and the confidence of the maximum cover has the lowest confidence.

After monosome gap filling, Falign applies the alignment selection method to the alignments selected from the chromosome. If the selection method returns a perfect cover from this chromosome (Figure 2E), it is highly likely that all fragments of this read exist on this chromosome. In this case, Falign outputs the perfect cover as the mapping results and terminates the aligning process for this read. If no perfect cover is found in this chromosome, Falign proceeds to the next chromosome containing the next highest-scoring candidate. These processes, including monosome candidate extension, gap filling and alignment selection, are performed on at most three chromosomes since the fragments of over 90% of Pore-C reads are spread across no more than three chromosomes.

### Polysomy gap filling and alignment selection

If no perfect cover is found from a single chromosome after processing all three chromosomes, the read may contain fragments from multiple chromosomes, and some fragments may not be sampled from these three chromosomes. Therefore, Falign conducts polysomy gap filling and alignment selection (Figure 2G and Methods). Falign first identifies the missing regions not covered by any alignments from the three chromosomes. Then, Falign collects k-mer chains that do not belong to those three chromosomes and sorts them by score. For each k-mer chain (starting with the one having the highest score), if over 80% of its coverage range is in the missing region, Falign performs the same extension procedure as that performed for long fragments and closes the missing region covered with the alignment. After gap filling, Falign performs alignment selection using all alignments. The returned alignment set, which can be either a perfect cover, a complete cover, or an incomplete cover, is output as the mapping result.

### Performance of Falign on simulated Pore-C datasets

We first evaluated Falign on three simulated Pore-C datasets (Figure 1, Methods and Supplementary Note 2). We compare Falign with bwa, NGMLR and minimap2 in terms of speed, accuracy, sensitivity and percentage of mapping (Methods and Supplementary Note 3). As shown in Table 1, Falign is 13.0-16.9 times faster than bwa and 1.5-6.5 times faster than NGMLR and minimap2 on the three datasets. The mapping results of Falign have no overlapped fragments, which indicates Falign has solved the fragment boundary problem. Falign has much higher recalls (93.45-95.59%) compared to the recalls of bwa (61.30-66.65%), NGMLR (48.50-57.03%), and minimap2 (77.79-84.38%), while all four aligners have comparable precisions.

We then analyzed the confident fragments extracted from their alignment results. A fragment is called confident if at least one of its alignment offsets is located at a restriction enzyme site. It is important that the fragment identified by the aligner is confident, as this can improve the reliability of the decomposition of a Pore-C read. The identification of confident fragments has been used for removing untrusted Hi-C alignments. For example, in the scaffolding tool LACHESIS^11^, Hi-C reads are treated as artifactual and excluded from subsequent analysis if they do not align within 500 bp of a restriction enzyme site. As shown in Table 1 and Figure 1G, over 99.59% of the fragments output by Falign are confident, significantly higher than the proportions output by bwa (71.21-88.99%), NGMLR (69.47-88.02%), and minimap2 (72.64-90.18%). The high percentage of confident fragments further demonstrates that Falign can properly identify the fragment boundaries.

The percentages of completely mapped reads (PCMRs) of Falign were 95.28-98.26%, which were significantly higher than those of minimap2 (8.16-11.73%), bwa (21.79-26.25%) and NGMLR (14.15-17.79%). The PCMR indicates the utilization of Pore-C data. The higher the PCMR is, the higher the percentage of Pore-C reads that have been utilized, and the more pairwise contacts that can be obtained. For example, the numbers of correct pairwise contacts decomposed from the mapping results of Falign are 30.17%, 27.96%, and 44.09% greater than those from the mapping results of minimap2 on the *H. sapiens, A. thaliana* and *D. melanogaster* datasets, respectively.

We next examine the effectiveness of gap filling in Falign. We implement another alignment method called Falign_ngf, which works in the same way as Falign except that neither monosome gap filling nor polysomy gap filling is performed in Falign_ngf. Falign_ngf was released together with Falign. As shown in Table 1, the sensitivities of Falign are 3.54-5.86% higher than those of Falign_ngf, and the PCMRs of Falign are 7.47-12.04% higher than those of Falign_ngf. As a result, Falign recovered 11.01-17.04% more correct pairwise contacts than Falign_ngf did. The results show that gap filling is effective in improving data efficiency.

To validate the generality of Falign to different restriction enzymes, we also generated simulated datasets using two other restriction enzymes: HindIII and NlaIII. Information about these datasets is listed in Supplementary Table 1. We also compare four aligners on these datasets. The results are summarized in Supplementary Table 2, from which we can draw the same conclusions as above.

### Performance of Falign on Pore-C datasets

We next evaluate Falign using three Pore-C datasets of *H. sapiens, A. thaliana* and *D. melanogaster* produced in-house using the DpnII restriction enzyme (Supplementary Table 3, Methods and Supplementary Notes 4-8). The error rates of the *H. sapiens* and *A. thaliana* datasets were distributed between 2-10% and were 5% on average (Supplementary Figure 5B). The quality of the *D. melanogaster* dataset was worse than that of the other two, the error rates of which were distributed between 5-20% and were 8% on average.

We then ran Falign, bwa, NGMLR and minimap2 on these datasets, and the results are summarized in Table 2. The four aligners had similar results to those obtained from the simulated datasets. bwa was still much slower than the other three aligners. Falign was 1.62-3.33 times faster than NGMLR and minimap2. There were no overlapping fragments or overlapping reads in the Falign mapping results, while there were 21.85%-47.32% overlapping fragments and 34.15%-75.66% overlapping reads in the bwa, NGMLR and minimap2 mapping results. The PCMRs in the mapping results of Falign were 89.60%, 87.88% and 72.57% on the three datasets, which were significantly higher than those in the mapping results of bwa (21.19%, 37.16% and 20.56%), NGMLR (16.90%, 32.28% and 31.02%) and minimap2 (10.93%, 21.91% and 18.54%). Benefiting from the significantly higher PCMRs, the numbers of pairwise contacts generated by Falign were 68.19%, 78.28% and 96.82% more than those generated by minimap2; 104.06%, 75.85% and 75.58% more than those generated by bwa; and 221.68%, 383.02% and 555.57% more than those generated by NGMLR on the three datasets. We also found that gap filling helped improve data utilization in Pore-C datasets. The PCMRs of Falign were 6.60%, 0.56% and 1.30% higher than those of Falign_ngf on *H. sapiens, A. thaliana and D. melanogaster*, respectively. Consequently, Falign generated 17.98%, 0.33% and 10.42% more pairwise contacts than Falign_ngf.

**Table 2.** Performance on Pore-C datasets. The datasets are generated using the DpnII restriction enzyme. The contacts are the pairwise contacts decomposed from the mapping results.

| Dataset | Aligner | Size (G)/<br>Time (h) | Fragment (%)<br>Confident/<br>Overlapping | Read (%)<br>Complete/<br>Overlapping | AFPR | Contacts (M) |
| --- | --- | --- | --- | --- | --- | --- |
| H. sapiens | bwa | 21.66/82.92 | 83.07/25.02 | 21.19/49.23 | 5.25 | 49.80 |
|  | NGMLR | 21.66/14.63 | 81.18/44.10 | 16.90/63.52 | 4.54 | 31.59 |
|  | minimap2 | 21.66/11.91 | 85.56/47.32 | 10.93/75.66 | 5.85 | 60.42 |
|  | Falign | 21.66/5.00 | 98.06/0.00 | 89.60/0.00 | 7.01 | 101.62 |
|  | Falign_ngf | 21.66/3.30 | 99.13/0.00 | 83.00/0.00 | 6.47 | 86.13 |
| A. thaliana | bwa | 11.38/9.41 | 78.92/21.85 | 37.16/34.15 | 3.65 | 19.09 |
|  | NGMLR | 11.38/2.50 | 73.38/39.59 | 32.38/47.19 | 2.93 | 6.95 |
|  | minimap2 | 11.38/2.03 | 81.95/43.79 | 21.91/60.38 | 3.84 | 18.83 |
|  | Falign | 11.38/1.25 | 98.56/0.00 | 87.88/0.00 | 4.14 | 33.57 |
|  | Falign_ngf | 11.38/1.10 | 98.58/0.00 | 87.32/0.00 | 4.13 | 33.46 |
| D. melanogaster | bwa | 29.96/34.47 | 79.65/23.94 | 20.56/40.30 | 4.02 | 55.97 |
|  | NGMLR | 29.96/6.33 | 87.63/33.33 | 31.02/39.89 | 2.84 | 14.99 |
|  | minimap2 | 29.96/5.68 | 89.57/37.86 | 18.54/54.38 | 3.95 | 49.93 |
|  | Falign | 29.96/1.90 | 98.36/0.00 | 72.57/0.00 | 4.79 | 98.27 |
|  | Falign_ngf | 29.96/1.51 | 98.30/0.00 | 71.27/0.00 | 4.57 | 89.00 |

Then, we analyzed Falign’s mapping results for the three datasets (Supplementary Figure 5, Methods and Supplementary Note 9). The three datasets have roughly the same distributions of fragment length. The average lengths of fragments in the three datasets are 919, 818 and 631, with the length of the majority of fragments being 200-1200 bp (Supplementary Figure 5A). A majority of the *H. sapiens* Pore-C reads contain 3-20 fragments, most of the *A. thaliana* reads contain 1-10 fragments, and most of the *D. melanogaster* reads contain 3-10 fragments (Supplementary Figure 5C). The orders of the *H. sapiens* Pore-C reads are much higher than those of the *A. thaliana* and *D. melanogaster* Pore-C reads, which is consistent with the fact that the *H. sapiens* Pore-C reads are almost twice as long as the *A. thaliana* and *D. melanogaster* Pore-C reads (Supplementary Table 3). In all three datasets, the fragments of over 50% are spread on a single chromosome, and the proportion of Pore-C reads decreases as the number of chromosomes on which their fragments are spread increases (Supplementary Figure 5D). The characteristics of Pore-C reads are close to those of the simulated datasets (Supplementary Figure 1).

### Performance of Falign on other long noisy 3C datasets

To verify that Falign also works well on other long noisy 3C datasets, we collected four publicly released human datasets (Supplementary Table 3): one GM12878 Pore-C dataset generated with NlaIII^17^, one GM24385 Pore-C dataset generated with HindIII^17^, one K-562 C-walks dataset generated with DpnII^23^, and one HelaS3 MC-3C dataset generated with DpnII^24^. The GM12878 and GM24385 datasets were sequenced by the ONT platform. The K562 and HeLaS3 datasets were sequenced with the PacBio platform^27^. We ran the four aligners on these datasets. As shown in Table 3, Falign still outperformed bwa, NGMLR and minimap2. There were no overlapping fragments or overlapping reads in the Falign results, while there were still significant percentages of overlapping fragments and overlapping reads in the bwa, NGMLR and minimap2 results. Falign still achieved higher data utilization than the other three aligners. The PCMRs of Falign were significantly higher than those of the other three aligners. In particular, on GM12878 and GM24385, Falign generated 103.24% and 20.24% more contacts than BWA, generated 2763.13% and 154.01% more contacts than NGMLR, and generated 49.40% and 112.54% more contacts than minimap2. The GM24385 data were generated using a six-base restriction enzyme (Hindlll). Therefore, the lengths of the fragments of its reads (average 4k bp) were significantly longer than those of the reads generated using four-base restriction enzymes. This caused the breakage of fragments during nanopore sequencing library construction, which led to fragments without restriction enzyme sites. Thus, all four tools reported low-confidence fragment rates (Table 3). However, Falign still reported a 90% read complete mapping rate, which was much higher than those reported by the other three tools. Overall, the experimental results show that Falign has an advantage in maximizing the data utilization of long noisy 3C data.

**Table 3.**
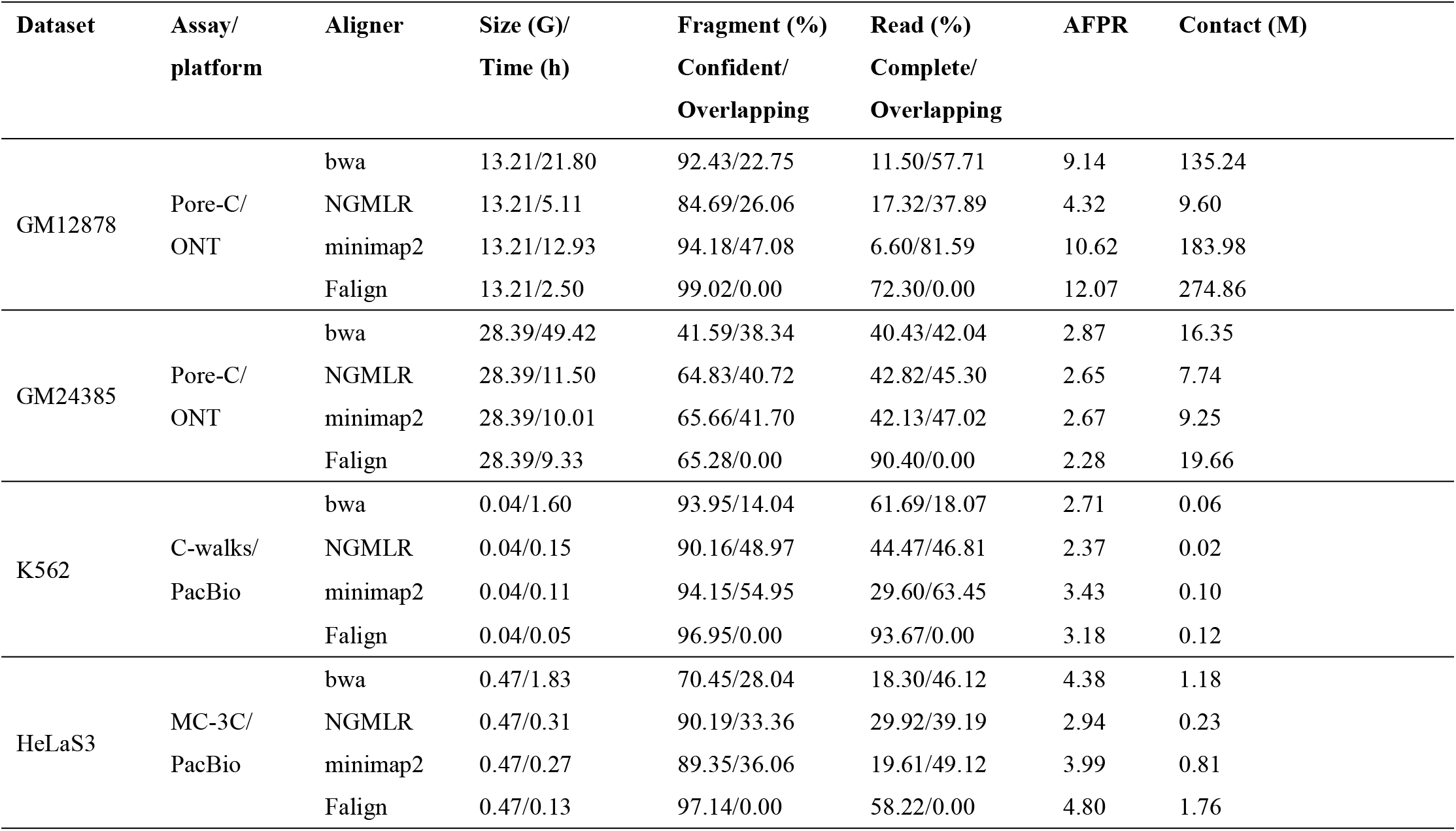
Performance on other 3C datasets.

## Discussion

FLNRs, such as Pore-C and C-walk, contain multiple fragments of varied length, which are separated by restriction enzyme motifs. In this paper, we demonstrated that the mainstream alignment tools have a low mapping rate for short fragments and find incorrect fragment boundaries when they are applied to Pore-C reads, which significantly affects the effective utilization of those reads for downstream studies. Several reasons lead to this low performance by existing alignment tools. First, none of the current alignment tools consider the fragmented nature of Pore-C reads. These tools do not stop at the boundaries of fragments during local alignment. Their local alignments can significantly exceed the fragment boundary (>50 bp), especially when aligning long fragments. This may lead to the loss of alignment to nearby short fragments. Second, the high error rates of reads means that alignment tools designed for next-generation sequencing (NGS), such as bwa, which cannot align because of the high-error region in the reads, miss a significant number of fragments. Third, the varied length of fragments means that alignment tools designed for long noisy reads, such as minimap2 and NGMLR, have difficulty selecting the parameter by which to align both long fragments and short fragments. The default parameters of these alignment tools tend to select long fragments and then miss the short fragments. However, if we chose lenient parameters, the computational cost would increase dramatically.

In this study, we develop Falign, a sequence alignment method that is adapted to the nature of fragmented long noisy reads. Falign adopts a two-phase approach. In the first phase, Falign finds the alignments for long fragments. It finds the alignments for short fragments in the second phase. This two-phase approach allows Falign to efficiently align both long fragments and short fragments. During the alignment of the fragments, Falign uses the restriction enzyme sites on the reference genome as boundaries, which avoids the problem that many fragment boundaries are destroyed because of sequencing errors or the differences between the underlying genome and the reference genome, such as SNPs. To effectively recover the alignments of Pore-C reads with multiple fragments and interchromosomal fragments, we employ a multiple-stage searching mechanism. If we find a solution with high confidence on a single chromosome (perfect cover), we stop searching and output the results. Otherwise, we extend all the seed chromosome candidates and choose the most plausible solution. The experimental results on simulated datasets show that Falign can effectively and completely recover the constructs of Pore-C reads and the sampled loci of the fragments. We also show that Falign performs equally well on Pore-C datasets generated with different restriction enzymes. The experimental results on Pore-C datasets show that Falign achieves significantly higher data utilization than the mainstream sequence aligners. We also demonstrate that Falign performs well on the other 3C datasets. Taken together, Falign can be a useful tool for fragmented long noisy reads.

## Data sources

We used nine simulated datasets and seven experimental datasets to evaluate the performance of the four aligners. Among these datasets, the Pore-C datasets of *H. sapiens, A. thaliana*, and *D. melanogaster* were generated using our in-house sequencing method (Supplementary Note 4-8), and nine simulated datasets were generated by our methods (Supplementary Note 1). The in-house simulated and experimental datasets is available from BIG (the Genome Sequence Archive in the BIG Data Center, Beijing Institute of Genomics, http://gsa.big.ac.cn) under Project Accession No. PRJCA012372.

## Accession codes

All processed files for the alignment and analysis code used in this study are available in the Supplementary Code, and all source code for Falign is available from https://github.com/xiaochuanle/Falign.

## ACKNOWLEDGMENTS

We thank all those who generated and freely released the data analyzed in our present study. This study was funded in part by the National Key R&D Program of China (2022YFF1201900), the National Natural Science Foundation of China (32270713, 31871326 and 91953122). We thank the Local Innovative and Research Teams Project of Guangdong Pearl River Talents Program (2017BT01S138) and the CAMS Innovation Fund for Medical Sciences (2019-I2M-5-005). This work was supported in part by grants from the U.S. National Institute of Food and Agriculture (NIFA; grant number 2017-70016-26051) and the U.S. National Science Foundation (NSF; grant numbers ABI-1759856, MRI-2018069, MTM2-2025541) to F.L.

## AUTHOR CONTRIBUTIONS

C.L.X. and F.L. conceived and designed this project. Y.C. and C.L.X. conceived, designed, and implemented the alignment algorithm. Y.C. integrated all the programs into the Falign pipeline and provided documentation. Z.B.L., J.Y.Z. and Y.C. ran analyzed genome assemblies and analyzed the performance of the algorithms developed in this study. L.J.N. constructed the Three Pore-C sequencing library. F.L., C.Y., C.H.H., Y.Z.L. and C.L.X. wrote the manuscript. All authors have read and approved the final version of this manuscript.

## COMPETING FINANCIAL INTERESTS

The authors have no competing financial interests to declare.

## Notes

https://github.com/xiaochuanle/Falign

### Competing Interest Statement

The authors have declared no competing interest.

## References

1. Stadhouders, R., Filion, G. J. & Graf, T. Transcription factors and 3D genome conformation in cell-fate decisions. Nature. 569, 345–354 (2019).

2. Dekker, J., Rippe, K., Dekker, M. & Kleckner, N. Capturing chromosome conformation. Science. 295, 1306–1311 (2002).

3. Simonis, M. et al. Nuclear organization of active and inactive chromatin domains uncovered by chromosome conformation capture-on-chip (4C). Nat. Genet.38, 1348–1354 (2006).

4. Zhao, Z. et al. Circular chromosome conformation capture (4C) uncovers extensive networks of epigenetically regulated intra- and interchromosomal interactions. Nat. Genet.38, 1341–1347 (2006).

5. Dostie, J. et al. Chromosome Conformation Capture Carbon Copy (5C): a massively parallel solution for mapping interactions between genomic elements. Genome Res.16, 1299–1309 (2006).

6. Fullwood, M. J. et al. An oestrogen-receptor-alpha-bound human chromatin interactome. Nature. 462, 58–64 (2009).

7. Lieberman-Aiden, E. et al. Comprehensive mapping of long-range interactions reveals folding principles of the human genome. Science. 326, 289–293 (2009).

8. Quinodoz, S. A. et al. Higher-Order Inter-chromosomal Hubs Shape 3D Genome Organization in the Nucleus. Cell. 174, 744–757 (2018).

9. Beagrie, R. A. et al. Complex multi-enhancer contacts captured by genome architecture mapping. Nature. 543, 519–524 (2017).

10. Ay, F. & Noble, W. S. Analysis methods for studying the 3D architecture of the genome. Genome Biol. 16, 183 (2015).

11. Burton, J. N. et al. Chromosome-scale scaffolding of de novo genome assemblies based on chromatin interactions. Nat. Biotechnol.31, 1119–1125 (2013).

12. Ghurye, J. et al. Integrating Hi-C links with assembly graphs for chromosome-scale assembly. PLoS Comput Biol. 15, e1007273 (2019).

13. Garg, S. et al. Chromosome-scale, haplotype-resolved assembly of human genomes. Nat. Biotechnol.39, 309–312 (2021).

14. Zhang, Y. et al. Chromatin connectivity maps reveal dynamic promoter-enhancer long-range associations. Nature. 504, 306–310 (2013).

15. Sanyal, A., Lajoie, B. R., Jain, G. & Dekker, J. The long-range interaction landscape of gene promoters. Nature. 489, 109–113 (2012).

16. Pope, B. D. et al. Topologically associating domains are stable units of replication-timing regulation. Nature. 515, 402–405 (2014).

17. Deshpande, A. S. et al. Identifying synergistic high-order 3D chromatin conformations from genome-scale nanopore concatemer sequencing. Nat. Biotechnol., (2022).

18. Zhong, J., Niu, L., Luo, F., Hou, C. & Xiao, C. Single-allele topology analysis with in situ HiPore-C reveals higher-order 3D genome folding principles., (2022).

19. Li, H. Aligning sequence reads, clone sequences and assembly contigs with BWA-MEM.; 2013. pp. 1303–3997.

20. Sedlazeck, F. J. et al. Accurate detection of complex structural variations using single-molecule sequencing. Nat. Methods. 15, 461–468 (2018).

21. Li, H. Minimap2: pairwise alignment for nucleotide sequences. Bioinformatics. 34, 3094–3100 (2018).

22. Xiao, C. L. et al. MECAT: fast mapping, error correction, and de novo assembly for single-molecule sequencing reads. Nat. Methods. 14, 1072–1074 (2017).

23. Olivares-Chauvet, P. et al. Capturing pairwise and multi-way chromosomal conformations using chromosomal walks. Nature. 540, 296–300 (2016).

24. Tavares-Cadete, F., Norouzi, D., Dekker, B., Liu, Y. & Dekker, J. Multi-contact 3C reveals that the human genome during interphase is largely not entangled. Nat. Struct. Mol. Biol.27, 1105–1114 (2020).

25. Li, Y. et al. DeepSimulator: a deep simulator for Nanopore sequencing. Bioinformatics. 34, 2899–2908 (2018).

26. Sosic, M. & Sikic, M. Edlib: a C/C ++ library for fast, exact sequence alignment using edit distance. Bioinformatics. 33, 1394–1395 (2017).

27. Korlach, J. et al. Real-time DNA sequencing from single polymerase molecules. Methods Enzymol. 472, 431–455 (2010).

