## Supplementary Figures for "Falign: An effective alignment tool for long noisy 3C data"

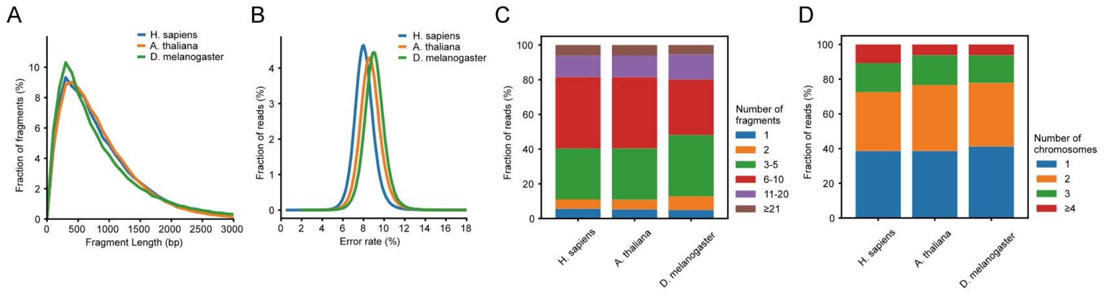


Supplementary Figure 1. Characteristics of simulated Pore-C datasets. (A) Distributions of fragment length. (B) Sequencing error rate distributions. (C) Distributions of Pore-C read orders. (D) Distributions of chromosome Pore-C read fragment spreading.

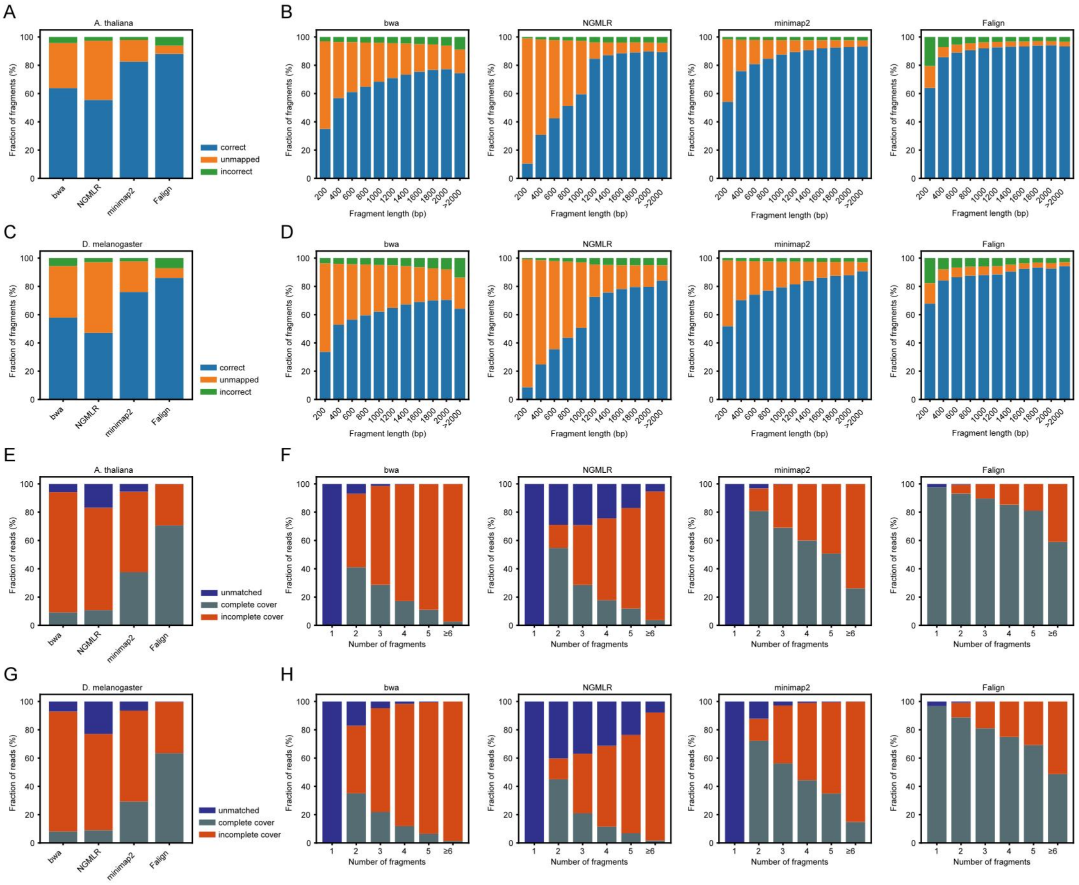


Supplementary Figure 2. Statistics of the mapping results of BWA, NGMLR, minimap2 and Falign on *A. thaliana* and *D. melanogaster* simulated Pore-C datasets. (A-D) Statistics of the mapping results in terms of fragments. (E-H) Statistics of the mapping results in terms of reads.

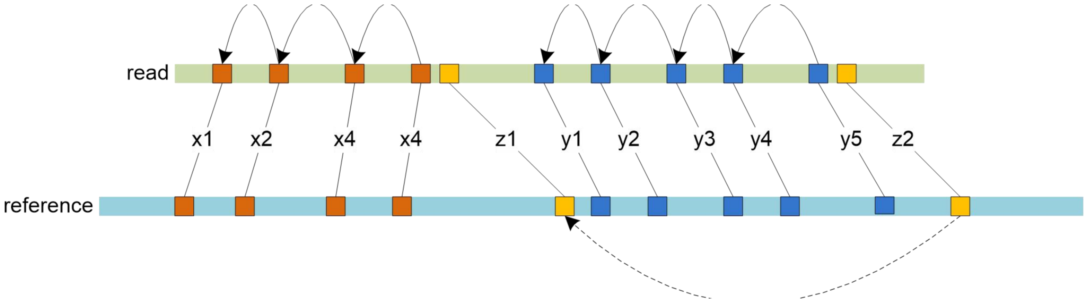


Supplementary Figure 3. Two $k$-mer chains (brown and blue) constructed by local chain scoring. The solid arrows are generated by backtracking using the local chain scoring formula (see Methods). The correlation (shown by the dashed arrow) of two remote $k$-mer matches is noninformative, although they support each other by DDF scoring because there exists no $k$-mer chain connecting them.

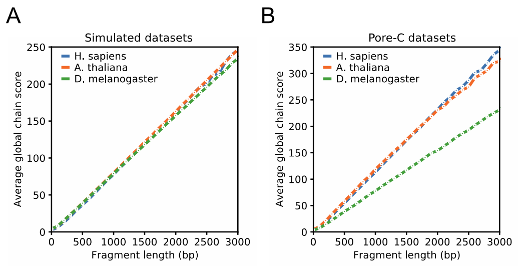


Supplementary Figure 4. The global chain scores of fragment candidates grow linearly on both simulated Pore-C reads (A, Table 1) and actual Pore-C reads (B, Table 2). For each dataset, we sequentially extracted 100,000 reads and then conducted statistical analysis of their mapping results.

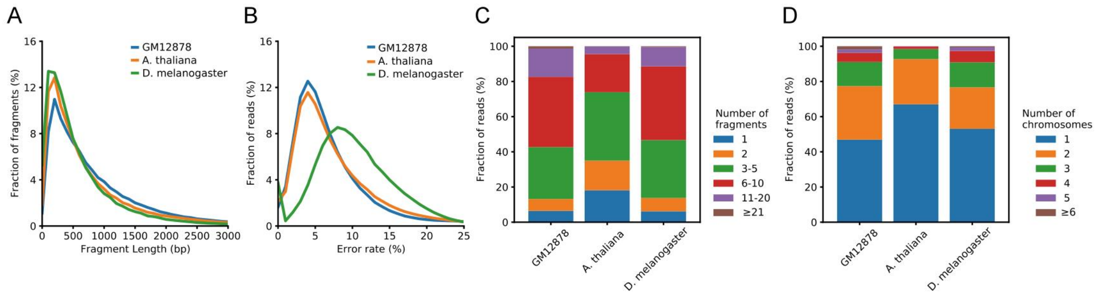


Supplementary Figure 5. Characteristics of Pore-C datasets. (A) Distributions of fragment lengths. (B) Distributions of sequencing error rates. (C) Distributions of Pore-C read orders. (D) Distributions of chromosome Pore-C read fragment spreading.
