## Supplementary Notes for "Falign: An effective alignment tool for long noisy 3C data"

### Supplementary Note 1: Construction of Simulated Pore-C Datasets

To evaluate the mapping accuracy of Falign, simulated *H. sapiens*, *D. melanogaster* and *A. thaliana* Pore-C datasets were constructed to provide known fragment sites and lengths. The features of Pore-C datasets, such as fragment number and chromosome source, were also considered. In brief, the prescribed reference genome was virtually cut by restriction enzyme sequences to collect virtual fragment sets. Then, randomly selected fragments were ligated to construct Perfect 3C datasets. Finally, the Nanopore sequence simulator transformed the perfect 3C reads into simulated Pore-C datasets. In total, we generated 9 Pore-C simulation datasets, each containing 2G bases. The details are listed in Supplementary Table 1 and Supplementary Figure 1.

**Perfect 3C dataset.** The latest versions of the *H. sapiens* (GRCh38), *D. melanogaster* (Release 6) and *A. thaliana* (TAIR10) reference genomes and three frequently used restriction enzymes, DpnII, HindIII and NlaIII, were used for simulation. The detailed information and download links are listed in Supplementary Table 1. Virtual restriction fragments were generated by directly cutting the genome by enzyme sequences. Realistically, folded DNA does not expose all restriction sites and generates fragments across several sites. To simulate this situation, one intact chromosome was randomly cut between 2-4 contiguous sites and generated virtual restriction fragments containing 1-3 basic fragments. With the fragment library constructed, the next step was proximity ligation. Considering the characteristics of Pore-C datasets, perfect 3C reads mainly consist of 3-20 fragments from one chromosome. To simulate this situation, we created a list of random numbers based on the proportion of fragment numbers in the Pore-C datasets. Based on the desired read number, we randomly selected seed fragments for proximity ligation and assigned fragment numbers for each read. Next, contact fragments of seed fragments were randomly selected based on the probability ratio of 4:2:1 from contiguous intrachromosomal sets (1/10 chromosome size), remote intrachromosomal sets and interchromosomal sets. At this point, the component of each read had been determined. DNA fragments were taken from both the forward and reverse strands. Based on the fragment number, we generated a list of random numbers 0 or 1. Fragments corresponding to 0 were from the forward strand and remained unchanged, while fragments corresponding to 1 were converted into their reverse-complement counterparts. Finally, all fragments of each read were linked together to generate Perfect 3C datasets. Fragment information was written in the header of the sequence for further analysis. Scripts for this procedure are provided in the Supplementary package.

**Nanopore sequencing simulation.** Nanopore sequencing signal simulation was conducted by DeepSimulator 1.5 with default parameters. DeepSimulator contains 3 modules, a Sequence Generator to produce DNA sequences from a reference genome (fasta), a Signal Generator to simulate electrical signals (fast5) of Nanopore sequencing and a Basecaller to generate simulated reads (fastq). To adapt to the Pore-C simulation, the Sequence Generator is modified to directly accept Perfect 3C datasets as input. Using the default parameters, signal simulation is performed by a context-independent pore model, and current signals are base called by the Guppy 3.1.5 HAC 'dna_r9.4.1_450bps_hac.cfg' model. The header of the simulated sequence is made up of 36 random characters in Nanopore format, and the matchup of the perfect 3C read and simulated read is provided in mapping.paf. For further analysis, one-to-one corresponding reads are preserved, and fragment information is added to the header of the sequence. At this point, the simulated Pore-C datasets have been constructed and used for evaluation. Scripts for this procedure are provided in the Supplementary package. All executive commands were written in deep_simulator.PoreC.sh to directly output simulated Pore-C datasets ready for use.

As an example, the following commands are used to generate the human DpnII Perfect 3C dataset:

```
reference_fasta=GCA_000001405.15_GRCh38_no_alt_plus_hs38d1_analysis_set.fna
enzyme_sequence=^GATC
main_chromosome=chr1
out_directory=Human_DpnII
python fragment_library.py -r ${reference_fasta} -e ${enzyme_sequence} -o ${out_directory}
python 3C_read_library.py -r ${reference_fasta} -e ${enzyme_sequence} -f
${out_directory}/fragments.parquet_0
```

The perfect 3C dataset is saved in:

```
${out_directory}/fragments.parquet_0.fasta
```

Then, the perfect 3C dataset is turned into the simulated Pore-C dataset:

```
./deep_simulator.PoreC.sh -i ${out_directory}/fragments.parquet_0.fasta -o ${out_directory}
```

The simulated Pore-C dataset is saved in:

```
${out_directory}/Csim.fastq
```

### Supplementary Note 2: Experimental Setup

The experiments in this paper are all performed on a computer equipped with 2 Intel Xeon E5-2678 CPUs (consisting of 48 2.50 GHz CPU threads) and 128 GB RAM. The versions of the aligners and their running options are listed in the following table.

| Aligner | Version | Running options |
|---|---|---|
| bwa | 0.7.17 | bwasw -b 5 -q 2 -r 1 -T 15 -z 10 |
| NGMLR | 0.2.7 | --subread-length 256 --max-segments 5 -x ont |
| minimap2 | 2.20 | -a -x map-ont -B 3 -O 2 -E 5 -k13 --secondary=no -Y |
| Falign | 0.0.1 | -num_threads 20 -outfmt paf |

The mapping results of bwa, NGMLR and minimap2 are generated in the following way. The aligners and corresponding running options are integrated into the Pore-C analysis pipeline by modifying the configure files. We first run them on the datasets with the corresponding options listed above. The `pore_c alignments assign-fragments` script in the Pore-C analysis pipeline is used to choose maximal cover alignments for each read. The results output by the `pore_c alignments assign-fragments` script are then used for evaluation. The output results of Falign are directly used for evaluation.

The following commands for the Pore-C pipeline carrying bwa, NGMLR and minimap2 are used to set the input files by configuring the files before running:

```
conda activate pore-c-snakemake
snakemake --use-conda -j 20
```

The following command for Falign is used to directly set the input files with command-line arguments:

```
Falign -num_threads 20 -outfmt paf ${enzyme_seq} ${reference_fasta} ${porec_fastq} -out map.paf
```

### Supplementary Note 3: Performance evaluation

The performance of the four aligners is evaluated in the following ways.
**Running time.** Using the default parameters and 20 CPU threads, Pore-C datasets are aligned by Falign and Pore-C pipeline carrying bwa, NGMLR and minimap2, and the running time is recorded by the time command of the Linux System Monitor.
The following command is used for the Pore-C pipeline carrying bwa, NGMLR and minimap2:

```
time snakemake --use-conda -j 20
```

The following command is used for Falign:

```
time Falign -num_threads 20 -outfmt paf ${enzyme_seq} ${reference_fasta} ${porec_fastq} -out map.paf
```

**Precision and sensitivity.** Simulated Pore-C datasets comprised of known fragments are used to evaluate the precision and sensitivity of the 4 aligners based on the distances between known sites and mapped sites. We define two fragments as having a correct pairwise relationship if the distance between the known site and the mapped site is less than 20 bp. The alignments obtained by each method fall into the following 3 cases: (1) correctly identifying the pairwise relationship (C); (2) identifying an incorrect pairwise relationship (I); and (3) failing to find the pairwise relationship (U). Thus, the precision of the aligner is defined as C/(C+I), and the sensitivity of the aligner is defined as C/(C+U). Moreover, the cumulative length of the correct alignments (C) is calculated.

The more fragments that are correctly recalled, the higher the completeness of the reads. In terms of Pore-C reads, we define a read as having complete coverage if all known fragments are recalled, having incomplete coverage if partial known fragments are recalled, and being unmatched if none of the known fragments are recalled.
**Confident fragments.** A Pore-C read comprises several fragments ligated by known restriction enzyme sites. A fragment is called confident if at least one of its alignment offsets is located at a restriction enzyme site. Each alignment is assigned to the nearest reference genome restriction fragment, and we define a confident end if the distance between the virtual site and mapped site is less than 20 bp. Thus, fragments are classified into 3 types: (1) double-ended confident; (2) single-ended confident; and (3) unconfident. The percentage of confident fragments indicates the reliability of the mapping positions of fragments. Moreover, the cumulative length of the confident alignments is calculated.
**Decomposition of Pore-C reads.** Ideally, the fragments of each Pore-C read are positioned end to end without gaps or overlaps; this is (1) a complete read. We further define (2) an incomplete read if large gaps (>50 bp) exist between adjacent fragments and (3) an overlapping read if large overlaps (>50 bp) exist. The percentage of complete reads and the percentage of overlapping fragments indicate the

completeness and reliability of the alignment results.

**Contacts.** A Pore-C read is called informative if it contains 2 or more fragments; the more fragments it contains, the higher the order of interaction and the more pairwise contacts it represents. The number of fragments is counted, and the virtual pairwise interactions between any 2 fragments are calculated by Combination Formula $C_N^2$. We use the average number of fragments per read and the total number of pairwise contacts to indicate the availability of the alignment results. For simulated Pore-C datasets, known pairwise contacts are constructed and compared to mapped pairwise contacts. Similar to fragments, we define the recall of pairwise contacts as the proportion of correct pairwise contacts (C) among the known pairwise contacts (C+U) and define the precision as the proportion of correct pairwise contacts among the mapped pairwise contacts (C+I). In terms of pairwise contacts, the recall and precision indicate the completeness and reconstruction of the genomic 3D architecture.

The `accuracy_stats.py` scripts for comparison between known fragments and alignments can only be used in simulated Pore-C datasets.

The following command is used for the Pore-C pipeline carrying bwa, NGMLR and minimap2:

```
python accuracy_stats.py -cq ${csim_fastq} -dir ${align_table}
```

The following command is used for Falign:

```
python accuracy_stats.py -cq ${csim_fastq} -paf map.paf
```

The `completeness_stats.py` scripts for the general statistics of confident fragments, decomposition of Pore-C reads and contacts are commonly used in Pore-C datasets and simulated Pore-C datasets.

The following command is used for the Pore-C pipeline carrying bwa, NGMLR and minimap2:

```
python completeness_stats.py -dir ${align_table}
```

The following command is used for Falign:

```
python completeness_stats.py -paf map.paf
```

### Supplementary Note 4: Cell Culture and Sequencing Material

A total of 7 datasets for 3 species (*H. sapiens*, *A. thaliana* and *D. melanogaster*) are used for training and testing our algorithms. Among these, 3 datasets were generated in our laboratory using a PromethION sequencer instrument (Oxford Nanopore Technologies, UK). Raw materials were cultured as described below:

**Human B lymphocytes GM12878 cell line culture.** The human B lymphocyte GM12878 cell line was purchased from Coriell Institute. Cells were cultured in T25 tissue culture flasks at a seeding density of 0.2 million viable cells/mL in 5 mL RPMI 1640 medium (Thermo Fisher, Cat# C11875500BT) supplemented with 15% fetal bovine serum (Gibco, Cat# 10100147), 2 mM L-glutamine and 1% penicillin/streptomycin. The cells were incubated at 37 °C under 5% carbon dioxide, and the medium was replaced every 3~4 days. Once the culture reached a plateau, approximately 1 million cells/mL, the cells were harvested for subsequent experiments or subcultured to maintain cell viability.

***D. melanogaster*** **OSC cell line culture.** A stable cell line of ovarian somatic cells (OSCs) was derived from the parental cell line fGS/OSS and stored at -80 °C. Cross and Sang's M3 (BF) medium was prepared from Shields and Sang M3 Insect Medium with L-glutamine (Sigma – Aldrich, Cat# S8398). Medium was supplemented with 10% fetal bovine serum, 10 mU/mL insulin and 10% fly extract. OSCs were cultured in 6-well plates at a seeding density of $10^4$ cells/mL and incubated at 25 °C. OSCs were trypsinized and counted for subsequent experiments or subculture.

***A. thaliana*** **Col-0 ecotype culture.** Wild-type Arabidopsis thaliana (L.) Heynh. Columbia-0 (Col-0) was used in this study. Seedlings were grown on half Murashige & Skoog (MS) plates at 22 °C with a 16 h/8 h light/dark regime. After 14 days, the aerial parts of these seedlings were harvested for subsequent experiments.

### Supplementary Note 5: Cross-linking

**GM12878.** Briefly, 10 million suspension cells were transferred to 50 mL centrifuge tubes and pelleted by centrifugation at 1000×g for 5 min. Cells were resuspended in 20 mL fresh serum-free medium and cross-linked with 1% formaldehyde for 10 min at room temperature (RT). Crosslinking was quenched by adding glycine to a final concentration of 125 mM and incubating for 5 min at RT. The cells were then washed three times in chilled 1× PBS and pelleted by centrifugation at 1000×g for 5 min at 4 °C between each wash. The fixed cell pellet was transferred to a 1.5 mL centrifuge tube, snap frozen with liquid nitrogen and stored at -80 °C.

***D. melanogaster* OSCs.** Approximately 10 million trypsinized cells were transferred to a 50 mL centrifuge tube and pelleted by centrifugation at 1500×g for 2 min. The cross-linking procedures were the same as those for GM12878 described above.

***A. thaliana*.** Up to 2.5 g aerial tissue was transferred to a 50 mL centrifuge tube and cross-linked with 1% formaldehyde in 15 mL freshly prepared nuclei isolation buffer (20 mM HEPES pH 8.0, 250 mM sucrose, 1 mM MgCl2, 5 mM KCl, 40% glycerol, 0.25% Triton X-100, 0.1 mM phenylmethanesulfonylfluoride, 0.1% β -mercaptoethanol) for 1 h at RT. Glycine was added to a final concentration of 125 mM and incubated under vacuum for 5 min at RT to quench cross-linking. The fixed tissue was then ground to a fine powder in liquid nitrogen and stored at -80 °C.

### Supplementary Note 6: DNA digestion and ligation

The Pore-C library was constructed as previously described. For the GM12878 and OSC cell lines, aliquots of 3 million crosslinked cells were lysed by incubation in chilled Hi-C lysis buffer (10 mM Tris-HCl pH 7.5, 10 mM NaCl, 0.2% NP-40) supplemented with 1× protease inhibitors (Roche, Cat# 11697498001) at 4 °C for 30 min with rotation. Nuclei were pelleted at 4 °C for 5 min at 1000 × g, and the supernatant was discarded. Pelleted nuclei were washed once with 500 µL of chilled Hi-C lysis buffer, resuspended in 50 µL 0.5% SDS and incubated at 62 °C for 10 min with no shaking or rotation. To quench SDS, 145 µL nuclease-free water and 50 µL 10% Triton X-100 were added, and the samples were incubated at 37 °C for 15 min with rotation.

For *A. thaliana*, the homogenate was resuspended in nuclei isolation buffer supplemented with 1× protease inhibitors and filtered through two layers of Miracloth. Nuclei were pelleted at 4 °C for 15 min at 3000 × g and washed twice with nuclei isolation buffer at 4 °C for 15 min at 1900 × g. For subsequent restriction enzyme digestion, nuclei were washed twice with 1.2 × NEB buffer 4 (NEB, Cat# B7004). Pelleted nuclei were resuspended in 500 µL 1.2 × NEB buffer 4 supplemented with 5 µL 20% SDS and incubated at 65 °C for 40 min and at 37 °C for 20 min with rotation. To quench SDS, 50 µL 20% Triton X-100 was added, and the samples were incubated at 37 °C for 1 h with rotation.

Chromatin was digested with 25 µL NEBuffer 3.1 (NEB, Cat# B7203) and 10 µL 10 U/µL DpnII restriction enzyme (NEB, Cat# R0543T) at 37 °C for 4 h with rotation. DpnII was then heat inactivated at 62 °C for 20 min with no shaking or rotation. For DNA ligation, 750 µL of ligation master mix was added: 100 µL 10× T4 DNA ligase buffer (NEB, Cat# B0202) with 10 mM ATP, 75 µL 10% Triton X-100, 3 µL 50 mg/mL BSA (Thermo Fisher, Cat# AM2616), 10 µL 400 U/µL T4 DNA Ligase (NEB, Cat# M0202), and 562 µL nuclease-free water. The reactions were then rotated at 16 °C for 4 h and RT for 1 h. Reverse cross-linking was performed by the addition of 10% SDS and 55 µl 20 mg/ml proteinase K and incubation at 63 °C overnight or for at least 4 h, followed by another addition of 65 µl 5 M NaCl and incubation at 68 °C for 2 h.

### Supplementary Note 7: DNA Purification

To isolate DNA, 500 µl phenol:chloroform:isoamyl alcohol (25:24:1) was added and mixed well. The mixture was transferred to a 2 mL MaXtract High Density tube (QIAGEN, Cat# 129056) to separate the aqueous phases, followed by the addition of 1 µl GlycoBlue (Thermo Fisher, Cat# AM9515), 100 µl 3 M sodium acetate, pH 5.2, and 850 µl isopropanol, and incubated at -80 °C for 1 h. Then, the precipitate was collected by centrifugation at 4 ℃ for 30 min at maximum speed, and the supernatant was discarded. The precipitate was washed twice in chilled 75% ethyl alcohol. After volatilization, the precipitate was resuspended in 170 µl Buffer EB (QIAGEN, Cat# 19086). Proteins were removed by adding 20 µl 10% SDS and 10 µl 20 mg/ml Proteinase K and incubating at 63 ℃ for 1 h. Another 100 µl phenol:chloroform:isoamyl alcohol (25:24:1) was added and mixed well, followed by centrifugation at 4 ℃ for 30 min at maximum speed, and the aqueous phase was collected. The aqueous phase was incubated in 20 µl 3M sodium acetate, pH 5.2, and 150 µl isopropanol at -80 °C for 1 h. The DNA was pelleted by centrifugation at 4 ℃ for 30 min at maximum speed and washed twice with chilled 75% ethyl alcohol. After volatilization, DNA was resuspended in 30 µl Buffer EB.

### Supplementary Note 8: Nanopore Sequencing and Base Calling.

Up to 3~4 μg of purified DNA per sample was qualified and used as input materials for nanopore library preparation. If needed, the 3C library was size selected using the PippinHT system (Sage Science, USA). Next, the sequencing libraries were constructed using a Ligation Sequencing Kit (Oxford Nanopore Technologies, Cat# SQK-LSK109). According to the manufacturer's instructions, end repair and dA tailing of DNA fragments were conducted with the NEBNext Ultra II End Repair/dA-tailing Module (NEB, Cat# E7546). Subsequently, 1D adaptor ligation was performed using the Quick Ligation Module (NEB, Cat# E6056). Finally, the prepared DNA library was measured by Qubit 4.0 Fluorometer (Invitrogen, USA). Approximately 700 ng of the DNA library was loaded and sequenced on a PromethION sequencer instrument (Oxford Nanopore Technologies, UK) at the Genome Center of Grandomics (Wuhan, China).

The raw reads were base called with Guppy 4.5.3 (Oxford Nanopore Technologies, UK) using the HAC (High-ACcuracy) 'dna_r9.4.1_450bps_hac_prom.cfg' model and default parameters. In total, 63 Gb of data were used in this study, generated from Pore-C libraries from the GM12878 cell line, *D. melanogaster* OSC cell line and *A. thaliana*. In addition to the intrinsic errors of nanopore sequencing, restriction enzyme processing could affect the quality of the library. To accommodate the Pore-C data, we adjust the base quality (Q) threshold to 7, while the default value is 9. Pass reads (Q>=7) were used for downstream analysis. The details are listed in Supplementary Table 3.

Basic statistics for Pore-C reads were conducted using Seqkit (2.0.0):

```
seqkit stats -a -j 20 ${porec_fastq}
```

### Supplementary Note 9: Characteristics of Pore-C Datasets

Unlike conventional nanopore reads, Pore-C reads consist of fragments that have different lengths and come from different chromosomes, and mapping is certainly more complicated. The alignment results of Pore-C are analyzed to show the characteristics of the Pore-C data, and the detailed results are shown in Supplementary Table 3 and Supplementary Figure 5.

**Fragment length.** Datasets from different species have similar mean fragment lengths: GM12878 has 919 bp, *A. thaliana* has 818 bp, and *D. melanogaster* has 631 bp. As shown in Supplementary Figure 5A, the 3 datasets possess similar fragment length distribution patterns and mainly range from 200 to 1000 bp, indicating that the fragment length is mainly correlated with the restriction enzymes rather than the species source or library length.

**Error rate.** The mapping consistency of each fragment is calculated using CIGAR string. The alignment match (M) base count is divided by the total base count, and the corresponding mapping error rate is equal to one minus the mapping consistency. Similarly, the distribution of the mapping error rate was separately analyzed for the 3 species datasets. The mean error rates of the GM12878 and *A. thaliana* datasets are 7.33% and 7.82%, respectively, while the *D. melanogaster* dataset possesses the highest error rate at 10.37%. The distribution of the mapping error rate of each dataset is shown in Supplementary Figure 5B. The difference in the mapping error rate between species could result from culture conditions and inconsistencies with the reference genome.

**Fragment number.** We calculated the mean fragment number per read by dividing the total fragment number by the mapped read number. Then, we statistically analyzed the fragment number distribution of each dataset. The mean fragment numbers of GM12878 and *D. melanogaster* are 6.96 and 6.29, respectively, which are higher than that of *A. thaliana* (4.21). In addition to the fact that datasets from different species have similar fragment lengths, the read length seems to be proportional to the fragment number. Most of the fragment numbers are in the range of 3~10, similar to the patterns of the 3 species datasets shown in Supplementary Figure 5C. Furthermore, GM12878 possesses a higher proportion of larger fragment number groups, indicating more high-order genomic interactions and a more complicated 3D genomic structure in humans.

**Chromosome source.** Chromosome conformation capture (3C) experiments can capture not only adjacent intrachromosomal contacts but also interchromosomal contacts. One Pore-C read is linked by fragments from different chromosome sites. We recorded the chromosome number of each dataset, and the mean chromosome numbers per read were 1.92 for GM12878, 1.42 for *A. thaliana* and 1.83 for *D. melanogaster*. The ratio of reads with different chromosome numbers is plotted in

Supplementary Figure 5D. As shown, the majority of reads (50~70%) consisted of adjacent fragments within the same chromosome, and the chromosome number was rarely more than 5. Importantly, more than 30% of the reads had fragments from different chromosomes, causing mapping difficulty in accurately decomposing the reads and completely aligning fragments to the reference genome.
