## Supplementary Tables for "Falign: An effective alignment tool for long noisy 3C data"

| Species | Ref. | Enzyme | Data size (Gb) | Read count | Mean read length | N50 (bp) | FL/FR |
|---|---|---|---|---|---|---|---|
| | | DpnII | 2.02 | 297,758 | 6,798 | 8,948 | 873/7.84 |
| H. sapiens | GRCh38 | HindIII | 2.01 | 190,329 | 10,557 | 17,607 | 4,709/2.26 |
| | | NlaII | 2.01 | 509,921 | 3,944 | 5,070 | 506/7.86 |
| | | DpnII | 2.02 | 306,236 | 6,582 | 8,546 | 844/7.85 |
| A. thaliana | TAIR10 | HindIII | 2.00 | 245,556 | 8,152 | 11,336 | 2,622/3.13 |
| | | NlaII | 2.00 | 319,185 | 6,271 | 8,096 | 806/7.84 |
| | | DpnII | 2.05 | 307,899 | 6,647 | 9,609 | 898/7.45 |
| D. melanogaster | Dmel R6 | HindIII | 2.03 | 136,601 | 14,853 | 20,629 | 4,816/3.10 |
| | | NlaII | 2.05 | 348,939 | 5,867 | 8,243 | 790/7.48 |

Supplementary Table 1. Statistics of simulated Pore-C datasets. FR: average number of fragments per read. FL: average fragment length.

| Dataset | Aligner | Size (G)/ Time (h) | Precision /Sensitivity (%) | Fragment (%) Confident/ Overlapping | Read (%) Complete/ Overlapping | AFPR map /raw | Contacts (M) map/correct/raw |
|---|---|---|---|---|---|---|---|
| **HindIII** | | | | | | | |
| H. sapiens | bwa | 2.01/3.71 | 53.21/41.90 | 77.91/41.29 | 38.82/46.80 | 1.78/2.26 | 0.49/0.14/0.55 |
| | NGMLR | 2.01/1.12 | 91.79/63.27 | 96.63/45.89 | 35.69/55.15 | 1.56/2.26 | 0.40/0.34/0.55 |
| | minimap2 | 2.01/0.52 | 97.07/69.05 | 97.73/52.70 | 30.35/62.42 | 1.61/2.26 | 0.42/0.40/0.55 |
| | Falign | 2.01/0.40 | 94.58/94.87 | 99.72/0.00 | 99.62/0.00 | 2.27/2.26 | 0.57/0.51/0.55 |
| | Falign_ngf | 2.01/0.37 | 95.06/94.15 | 99.87/0.00 | 99.27/0.00 | 2.24/2.26 | 0.52/0.49/0.55 |
| A. thaliana | bwa | 2.00/1.46 | 79.32/59.11 | 90.89/22.99 | 56.79/26.04 | 2.33/3.13 | 0.69/0.42/1.19 |
| | NGMLR | 2.00/0.49 | 92.74/73.87 | 96.84/45.47 | 36.08/54.06 | 2.49/3.13 | 0.80/0.70/1.19 |
| | minimap2 | 2.00/0.32 | 96.96/82.80 | 97.67/51.86 | 30.11/61.62 | 2.67/3.13 | 0.89/0.84/1.19 |
| | Falign | 2.00/0.13 | 96.44/96.13 | 99.96/0.00 | 99.43/0.00 | 3.12/3.13 | 1.18/1.11/1.19 |
| | Falign_ngf | 2.00/0.12 | 96.43/95.42 | 99.95/0.00 | 98.87/0.00 | 3.10/3.13 | 1.15/1.08/1.19 |
| D. melanogaster | bwa | 2.03/2.18 | 57.24/49.12 | 79.58/36.50 | 42.60/40.90 | 2.66/3.10 | 0.55/0.16/0.65 |
| | NGMLR | 2.03/0.53 | 86.54/68.53 | 95.12/44.92 | 33.73/54.10 | 2.46/3.10 | 0.44/0.33/0.65 |
| | minimap2 | 2.03/0.29 | 96.22/80.34 | 97.51/51.61 | 30.22/61.32 | 2.59/3.10 | 0.47/0.44/0.65 |
| | Falign | 2.03/0.25 | 94.51/94.32 | 99.91/0.00 | 99.36/0.00 | 3.10/3.10 | 0.65/0.59/0.65 |
| | Falign_ngf | 2.03/0.23 | 94.50/92.74 | 99.93/0.00 | 98.15/0.00 | 3.05/3.10 | 0.61/0.56/0.65 |
| **NlaIII** | | | | | | | |
| H. sapiens | bwa | 2.01/3.19 | 94.14/64.89 | 89.57/25.53 | 22.29/49.91 | 5.42/7.86 | 10.18/9.05/21.91 |
| | NGMLR | 2.01/0.90 | 94.87/21.26 | 86.87/32.74 | 20.23/44.17 | 1.76/7.86 | 1.75/1.59/21.91 |
| | minimap2 | 2.01/0.93 | 97.02/82.05 | 90.76/56.32 | 8.63/84.18 | 6.65/7.86 | 15.57/14.69/21.91 |
| | Falign | 2.01/0.33 | 94.05/93.34 | 99.69/0.00 | 96.37/0.00 | 7.80/7.86 | 21.53/18.90/21.91 |
| | Falign_ngf | 2.01/0.25 | 94.21/88.04 | 99.83/0.00 | 83.83/0.00 | 7.35/7.86 | 18.17/16.09/21.91 |
| A. thaliana | bwa | 2.00/1.35 | 94.02/67.31 | 71.75/25.39 | 22.81/50.92 | 5.61/7.84 | 6.93/6.16/13.69 |
| | NGMLR | 2.00/0.60 | 95.46/54.76 | 69.64/47.76 | 15.13/68.58 | 4.50/7.84 | 5.08/4.65/13.69 |
| | minimap2 | 2.00/0.57 | 97.52/84.94 | 73.07/57.14 | 8.58/85.57 | 6.83/7.84 | 10.33/9.84/13/69 |
| | Falign | 2.00/0.10 | 93.54/93.77 | 99.97/0.00 | 97.72/0.00 | 7.79/7.84 | 13.23/11.26/13.69 |
| | Falign_ngf | 2.00/0.10 | 95.88/92.02 | 99.97/0.00 | 90.18/0.00 | 7.52/7.84 | 12.22/11.20/13.69 |
| D. melanogaster | bwa | 2.05/1.70 | 91.74/63.42 | 84.80/24.00 | 26.93/45.44 | 5.17/7.48 | 6.82/5.78/14.40 |
| | NGMLR | 2.05/0.74 | 93.62/46.67 | 83.67/44.77 | 18.00/62.55 | 3.73/7.48 | 4.16/3.68/14.40 |
| | minimap2 | 2.05/0.65 | 96.61/79.96 | 86.75/54.04 | 11.59/79.42 | 6.19/7.48 | 9.87/9.25/14.40 |
| | Falign | 2.05/0.13 | 94.17/93.54 | 99.96/0.00 | 97.07/0.00 | 7.43/7.48 | 14.18/12.60/14.40 |
| | Falign_ngf | 2.05/0.11 | 94.59/88.21 | 99.97/0.00 | 86.64/0.00 | 6.98/7.48 | 12.09/10.94/14.40 |

Supplementary Table 2. Performance on simulated Pore-C datasets. The datasets were generated using the HindIII and NlaIII restriction enzymes.

| Platform | Method | Cell line (or species) | Enzyme | Data size (Gb) | Read count | Mean len. (bp) | N50 (bp) | Download datasets |
|---|---|---|---|---|---|---|---|---|
| ONT | Pore-C | GM12878 | DpnII | 21.66 | 3,336,569 | 6492 | 7564 | HRA003207 |
| | | A. thaliana | DpnII | 11.38 | 3,142,952 | 3620 | 4329 | CRA008434 |
| | | D. melanogaster | DpnII | 29.96 | 7,488,684 | 4001 | 4486 | CRA008433 |
| | Pore-C | GM12878 | NlaIII | 13.21 | 2,458,481 | 5373 | 7873 | NA12878.tar.gz |
| | | GM24385 | HindIII | 28.39 | 7,432,225 | 3820 | 6849 | HG002.tar.gz |
| PacBio | C-walks | K562 | DpnII | 0.04 | 21,771 | 1770 | 1987 | GSE77553 |
| | MC-3C | HeLaS3 | DpnII | 0.47 | 128,381 | 3685 | 4552 | GSE146945 |

Supplementary Table 3. Statistics of 3C datasets.
